# Linked networks reveal dual roles of insect dispersal and species sorting for bacterial communities in flowers

**DOI:** 10.1101/847376

**Authors:** Ash T. Zemenick, Rachel L. Vannette, Jay A. Rosenheim

## Abstract

Due to the difficulty of tracking microbial dispersal, it rarely possible to disentangle the relative importance of dispersal and species sorting for microbial community assembly. Here, we leverage a detailed multilevel network to examine drivers of bacterial community assembly within flowers. We show that plant species with similar visitor communities tend to have similar bacterial communities, and visitor identity to be more important than dispersal rate in structuring floral bacterial communities. However, plants occupied divergent positions in plant-insect and plant-microbe networks, suggesting an important role for species sorting. Taken together, our analyses suggest dispersal is important in determining similarity in microbial communities across plant species, but not as important in determining structural features of the floral bacterial network. A multilevel network approach thus allows us to address features of community assembly that cannot be considered when viewing networks as separate entities.

## Introduction

The strong but variable effects of microbial communities on ecological interactions and host fitness is now well-recognized (Friesen 2013, Sugio et al. 2015). However, it remains difficult to determine the relative influence of factors such as dispersal and species sorting influencing microbiome assembly, composition and function. This difficulty remains in no small part because it is difficult to track microbial dispersal in natural systems, and as a result, most study of microbial community assembly uses spatial distance as a proxy for dispersal (Venkataraman et al. 2015, Burns et al. 2016), or assumes global dispersal. However, it is clear that microbes can be dispersal limited (Peay et al. 2012). As a result, understanding the relative contribution of microbial dispersal compared to environmental factors has critical applications for understanding the transmission of not only pathogens but beneficial and commensal microbes, as well as the patterns underlying community assembly of complex microbiomes.

One way to assess the relative influences of dispersal and species sorting on community structure is through the use of coupled networks (Pilosof et al. 2017). In this framework, multiple linked networks are sampled, and analyzed together. Here, we use diagonally coupled networks (Pilosof et al. 2017) to compare structure between host-vector networks and host-microbe networks to examine potential effects of vector dispersal or species sorting on microbial community composition. Comparing network macro- and microstructure between linked networks can indicate processes underlying assembly.

The macrostructure of bipartite ecological networks can be nested or modular, which can differentially influence dispersal dynamics (Fig. 1). For example, a host-microbe network could be modular if microbes are highly specialized and experience strong species sorting, or, if their vectors are highly specialized and thereby limit dispersal to few host species. In contrast, nested host-bacteria networks may reflect weaker species sorting and dispersal limitation relative to modular networks. Macrostructural properties (e.g., nestedness, modularity) can not only inform the potential for microbial dispersal throughout a network (Tylianakis et al. 2010, Silk et al. 2017), but can be compared across networks to infer the relative importance of dispersal vs. species sorting, where significant correlations of network properties between linked host-vector and host-microbe networks would suggest the importance of dispersal.

**Figure 1.**
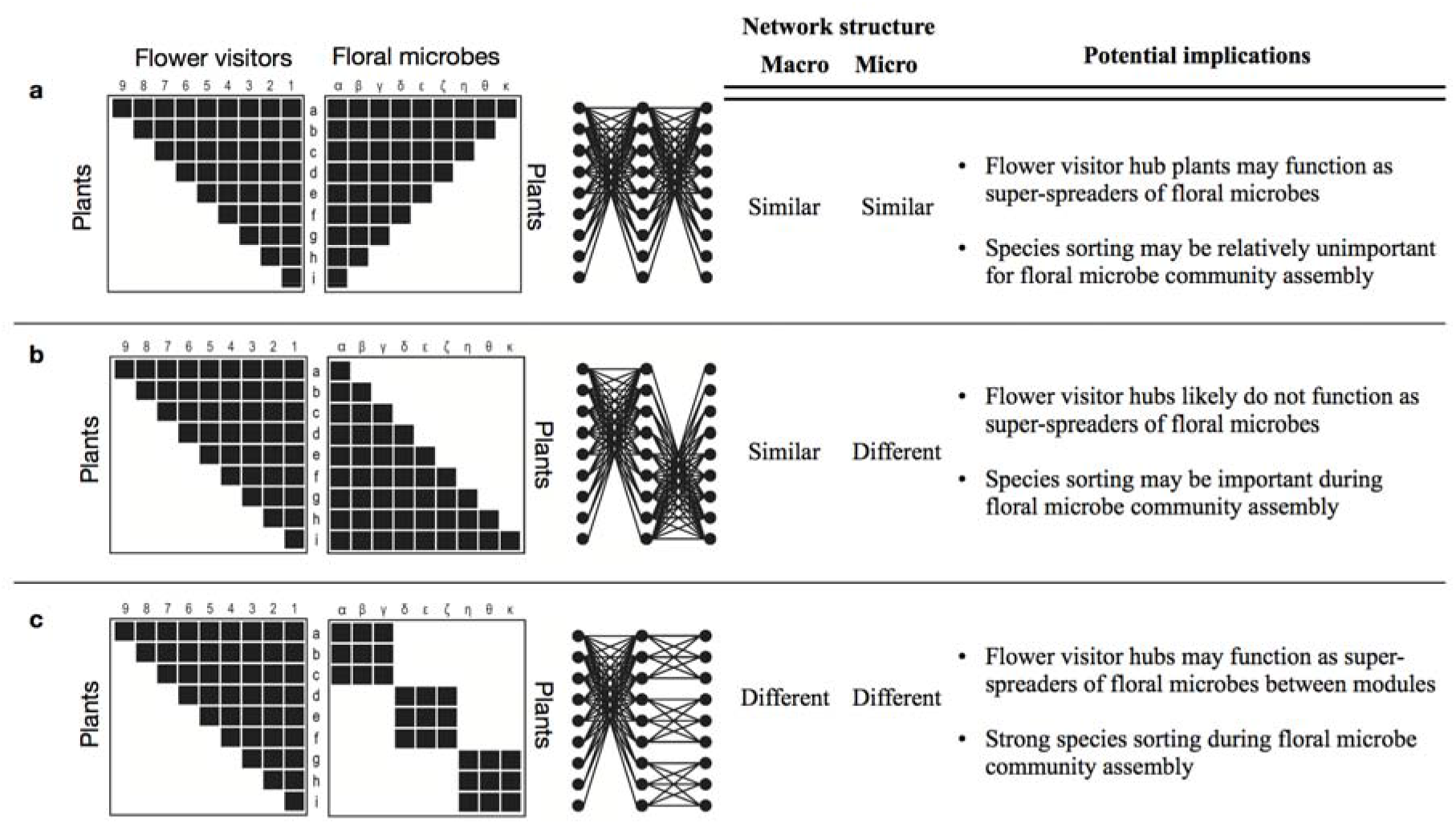
A conceptual framework of a multilevel plant – flower visitor – floral microbe network, where, for simplicity, each matrix is either perfectly nested or modular. Matrix rows represent plant species (a-i) and columns represent flower visitor (1-9) or floral microbe (⍰-κ) species. The figure is modified from Fontaine et al. (2011). If floral microbe networks share macrostructure with its associated flower visitor network, they may be linked in such a way that plants occupy similar (a) or dissimilar (b) positions within each network. If floral microbe communities are modular (c), then the relative position of plants in each network would depend on the structure of modules.

Network properties can also identify host species of particular importance for bacterial dispersal dynamics. By comparing the relative positions of species within networks and the specific identities of their interactions (i.e. network microstructure), we can identify hubs of interaction, which can disproportionately influence disease transmission in human and wildlife systems (Stein 2011, Silk et al. 2017). Hubs can amplify spread by interacting with many other individuals or species (i.e., species with high degree), link otherwise separated modules together (i.e., species with high betweenness centrality), or reduce the average path length connecting two species through shared interactors (i.e. species with high closeness centrality; Silk et al. 2017). If a host plant species is a primary hub of both the vector-host plant network and the host plant-bacteria network (i.e. the networks have similar microstructures), then this host species should have a very strong influence on metacommunity structure by facilitating high dispersal rates to other species in the community in an environment that is not very selective, making species sorting weak.

Floral microbial communities offer an ideal system in which to apply a multilevel network approach. First, floral microbial communities change over time (Shade et al. 2013, Aleklett et al. 2014) and influence plant and pollinator health and mediate plant-pollinator interactions (Ngugi and Scherm 2006, Vannette et al. 2013, Junker et al. 2014, McArt et al. 2014, Graystock et al. 2015, Rering et al. 2017). Flower-visiting insects (Ushio et al. 2015) and hummingbirds (Belisle et al. 2012) introduce microbial propagules to flowers. Yet, plant hosts may also exert selection on microbial establishment and growth through multiple mechanisms (Carter and Thornburg 2004, Huang et al. 2012, Block et al. 2019). Although floral microbe community assembly is dependent, in part, on dispersal by flower visitors, the study of floral microbe community assembly has to date been limited to single plant species or coarsely characterized communities of floral visitors, and remains yet to be fully considered in the context of the rich network of plant – flower visitor interactions.

Based on the conceptual framework of multilevel networks proposed by Fontaine et al. (2011), we outline three scenarios in which flower-visitor (host plant-vector) and host plant-bacteria networks could be linked, and what this structure suggests about dispersal and species sorting. If, like flower visitor networks (Bascompte et al. 2003), floral microbe networks are nested, the networks could be linked in two ways. Plant species may occupy similar positions in each network, suggesting that dispersal by flower visitors is a stronger force than species sorting in microbial community assembly (Fig. 1a). Alternatively, the networks could be aligned such that plants occupy different positions in each network, which could arise if plant traits filter microbial colonization (species sorting; Fig. 1b). If floral microbe networks are modular, and dispersal by visitors is more important than species sorting in floral microbe community assembly, then flower visitor hubs may be important in linking modules in the floral microbe network (Fig. 1c). Since visitors can be important vectors of floral microbes, we hypothesized that plant species should occupy similar positions in each network and that species with high diversity of visitors should have high diversity of microbes, functioning as hubs of both networks (Fig. 1a). To test this hypothesis, we constructed the first comprehensive snapshot of a host plant – flower visitor – floral microbe network to infer the importance of dispersal and species sorting in this metacommunity. We asked: (1) Can variation in bacterial species composition within and across plant species be explained by visitor frequency or visitor species composition (after accounting for host plant relatedness)? (2) Do flower visitor and floral bacterial networks have similar macrostructure (i.e., are both similarly nested or modular)? (3) Do flower visitor and floral bacterial networks have similar microstructure (i.e. are visitor hub plants also bacterial hub plants)?

We found that plant species with similar visitor communities tend to have similar bacterial communities, and that visitor identity may be more important than dispersal rate in structuring floral bacterial communities. While both the plant-bacteria and plant-visitor networks were significantly nested, plants occupied divergent positions in each network, suggesting an important role for species sorting. Taken together, it seems that dispersal is important in determining similarity in microbial communities across plant species, but not as important in determining structural features of the floral bacterial network.

## Materials and Methods

### Data collection

#### Study site

Flower visitor and floral microbe surveys were conducted in a high elevation wet meadow at the University of California’s Sagehen Creek Field Station, located within Tahoe National Forest (2400m, 39°25’11.52”N, 120°18’27.18”W). The meadow is dominated by herbaceous perennial flowering plants and is surrounded by a subalpine old growth pine-fir forest. The focal plant community included the 20 most abundant co-flowering plant species: *Ligusticum grayi* (Apiaceae), *Arnica mollis* (Asteraceae)*, Erigeron coulteri* (Asteraceae), *E. glacialis* (Asteraceae), *Senecio triangularis* (Asteraceae), *Lupinus polyphyllus* (Fabaceae), *Trifolium kingii* (Fabaceae), *Lilium parvum* (Liliacae)*, Triantha occidentalis* (Liliacae), *Veratrum californicum* (Melanthiaceae), *Platanthera dilatata* (Orchidaceae)*, P. sparsiflora* (Orchidaceae), *Castilleja miniata* (Orobanchaceae)*, Pedicularis groenlandica* (Orobanchaceae), *Mimulus guttatus* (Phrymaceae), *M. primuloides* (Phrymaceae), *Bistorta bistortoides* (Polygonaceae), *Aconitum columbianum* (Ranunculaceae), *Aquilegia formosa* (Ranunculaceae), and *Drymocallis lactea* (Roseaceae).

#### Flower visitor communities

During a ten-day window in July 2015, each of the 20 focal plant species was observed for all flower-visiting arthropods (regardless of their size or perceived pollination efficiency) during fourteen 30-minute time periods (7 hours of observation per species, 140 total observation hours). Observations for plant species were randomly assigned to time of day (08:00-17:00 h) and to 1 of 5 observers. On observation days, each species was observed once in the morning and once in the afternoon. The first ten minutes of each observation window focused on the observation and collection of micro-flower visitors using short focal length binoculars as a visual aid. The final twenty minutes were used to collect any flower visitors that could be seen with the unaided eye. Each flower-visiting arthropod observed during these time windows was collected and identified to the lowest taxonomic level possible by the authors and taxonomic experts (see Acknowledgements).

#### Floral microbial communities

At two time-points during the flower visitor observations, 15 flowers of each plant species were collected for floral microbe community assessment (30 flowers per species for a total of 600 flowers). Flowers of each species were chosen haphazardly such that they were representative of the whole study meadow and life stages of the flower. Flowers were placed in sterile plastic bags and frozen at −20°C then −80°C until processing.

Floral microbial DNA was isolated by placing five flowers per species into phosphate buffer saline (PBS), sonicating for 10 minutes, vortexing briefly, and then collecting resulting microbe + PBS solution by filtering through autoclaved cheesecloth (Shade et al. 2013). The solution was centrifuged at 3000 rpm for 10 min at 20°C, and the resulting pellet was used for extraction. Floral microbial DNA was extracted from each sample using the Qiagen DNeasy 96 Blood and Tissue Kit following the manufacturer protocols. Extracted DNA was sent to the Microbiome Resource Center (Halifax, Nova Scotia, Canada) for amplicon library preparation and MiSeq Illumina sequencing using ITS2 and 16S V4-V5 for fungi and bacteria respectively. Amplification of chloroplast DNA in 16S reactions was reduced with pPNA PCR blockers (GGCTCAACCCTGGACAG; PNA Bio, Newbury Park, CA).

Sequence data were quality filtered and grouped into operational taxonomic units (OTUs) using the UPARSE pipeline. First, low quality trailing bases were removed with sickle (Joshi and Fass 2011). Next, we merged read pairs with usearch version 5.1 (Edgar 2010), but due to the low merging success rate (32.6%), only forward reads were used in analyses (McFrederick et al. 2017). All sequences represented by only one read were removed, and sequences were clustered into OTUs with a 97% similarity cutoff with usearch (Edgar 2010). Chimeras were removed using *de novo* detection in usearch (Edgar 2010) and via referencing the Gold database (drive5.com/uchime/gold.fa). OTUs were then classified taxonomically with the Ribosomal Database Project (RDP) Naive Bayesian rRNA Classifier Version 2.11 (Wang et al. 2007). Bacterial taxonomical hierarchy was classified in reference to the RDP 16S rRNA training set 16, and fungi with the Warcup Fungal ITS trainset 2 (Deshpande et al. 2016). OTU and taxonomy tables were assembled and analyzed in R (R Core Team 2016) using the phyloseq package (McMurdie and Holmes 2013).

Few reads from the ITS primers were recovered, perhaps due to uneven or low fungal abundance, so only 16S amplicon data were utilized for analyses. From these amplicons, Cyanobacteria and Chloroplast OTUs were removed. No sequences were recovered from DNA extraction controls. In total, 2,927 bacterial OTUs were detected. To remove bias of uneven sequence depth across samples, all samples were rarified to an even depth of 4,206 reads to match the sample with the lowest number of reads. Sampling curves indicated that this sampling depth captures OTU richness and Shannon diversity of samples (see Fig. S1 in Supporting Information). After rarefaction, 2,135 OTUs were represented in the dataset.

### Data analysis

All analyses were performed in R version 3.3.3 (R Core Team 2016).

#### Variation in bacterial communities within and across co-flowering plant species

To assess whether plant species host different bacterial communities, we compared multivariate community dispersion and species composition of floral bacteria across samples using the functions ‘betadispr’ and ‘adonis’ based on Bray-Curtis distances using the R package vegan (Oksanen et al. 2017). Random forest analysis was used to identify the bacterial OTUs that were most important in distinguishing the bacterial species composition among different plant species using randomForest (Liaw and Wiener 2002).

To examine if flower visitor and microbial communities were correlated at the plant species level, Jaccard and Bray-Curtis dissimilarity matrices were constructed using the “vegdist” function (Oksanen et al. 2017). A Mantel test was used to examine the correlations between matrices. To evaluate whether particular groups of flower visitors were more strongly associated with variation in floral bacterial community patterns, the analysis was repeated using dissimilarity matrices based on different subsets of the flower visitor community: bees (39 species), non-bee Hymenoptera (120 species), Diptera (95 species), and Coleoptera (35 species). See Fig. S2 for a breakdown of the visitor community across plant species.

Since a correlation between visitor and bacterial communities could be due to bacteria and visitors tracking similar plant traits, we constructed a plant phylogenetic distance matrix to serve as a proxy for similarity in functional plant traits (see Fig. S3). Relatedness of the focal plant species was determined by reconstructing a phylogeny with the Interactive Tree of Life (Letunic and Bork 2006), which was converted to a phylogenetic distance matrix using the package ape (Paradis et al. 2004). A partial Mantel test was used to test for a correlation between the flower visitor dissimilarity matrix and the floral bacteria dissimilarity matrix, while taking into account the plant phylogenetic dissimilarity matrix.

#### Flower visitor and floral bacterial network architecture

##### Network macrostructure

To compare the macrostructural properties of the flower visitor and floral bacteria networks, we assessed the degree to which each network was nested and modular. Nested networks have a core of generalist species that interact, and highly specialized species tend to interact with subsets of the generalist core; (Bascompte et al. 2003). Network nestedness was estimated using the “nested” function in bipartite (Dormann et al. 2008, 2009) with methods “NODF2,” which considers only the presence of interactions, and “weighted NODF,” which also accounts for interaction frequencies. Both metrics range from 0-100, where 100 indicates a perfectly nested network. Modular networks are more partitioned, and are composed of species groups that interact more strongly with each other than with other species in the community (Poisot 2013). Modularity was assessed using the “compart” and “computeModules” functions in bipartite. The first counts the number of distinct groups of plants and visitors or bacteria that interact only with each other, and not any other members in the community (“compartments”). The second modularity metric (“Modularity Q”) identifies modules in which “within-module interactions are more prevalent than between-module interactions,” but without requiring that modules are discrete compartments (Dormann and Strauss 2013). Modularity Q ranges from 0 (less modular) to 1 (more modular).

Because network size affects measures of nestedness and modularity (Nielsen and Bascompte 2007, Fründ et al. 2016), and because there were many more bacterial OTUs than species of flower visitors (2,135 vs. 289), the floral bacteria network was rarefied by sampling randomly without replacement for 289 bacterial OTUs. The 95% confidence intervals of the rarefied bacterial network (based on 1000 rarefied networks) were used to determine whether metrics calculated for the floral bacteria network differed significantly from the flower visitor network. We also report metrics quantified from the un-rarefied bacterial network for comparison.

##### Network microstructure

Next, we compared the position of plant species in the flower visitor and floral bacteria networks. For each plant species, we compared the proximity to the nested core in each network. We then assessed whether plant species that are hubs of the flower visitor network are also hubs of the floral bacteria network by comparing degree (richness of interaction partners) and weighted closeness centrality (a measure of how closely linked a species is to all other species in the network; Silk et al. 2017) for plants in terms of their interactions with flower visitors and associations with floral bacteria. Closeness centrality was used instead of betweenness centrality due to the low levels of modularity in each network. Proximity to nested core, degree, and weighted closeness centrality were computed in bipartite with the ‘specieslevel’ function and indices “nestedrank”, “degree”, and “closeness.” Linear models were used to assess whether indices were significantly correlated across network types. For computational efficiency, closeness centrality and proximity to the nested core for the floral bacteria network were quantified based on an interaction matrix between plant species and bacterial genera instead of OTUs. The 2,136 OTUs were represented by 644 genera.

## Results

### Variation in bacterial communities within and across co-flowering plant species

Plant species differed in floral bacteria species composition (Fig. 2; Fig. S4; perMANOVA R^2^ = 0.77, p = 0.001) and also differed in dispersion of bacterial species composition (F_19_ = 1.7, p = 0.048). Bacterial taxa that best distinguished plant species included representatives from the classes Actinobacteria, Alphaproteobacteria, Bacilli, Betaproteobacteria, and Gammaproteobacteria (Fig. S5, Table S1, Table S2).

**Figure 2.**
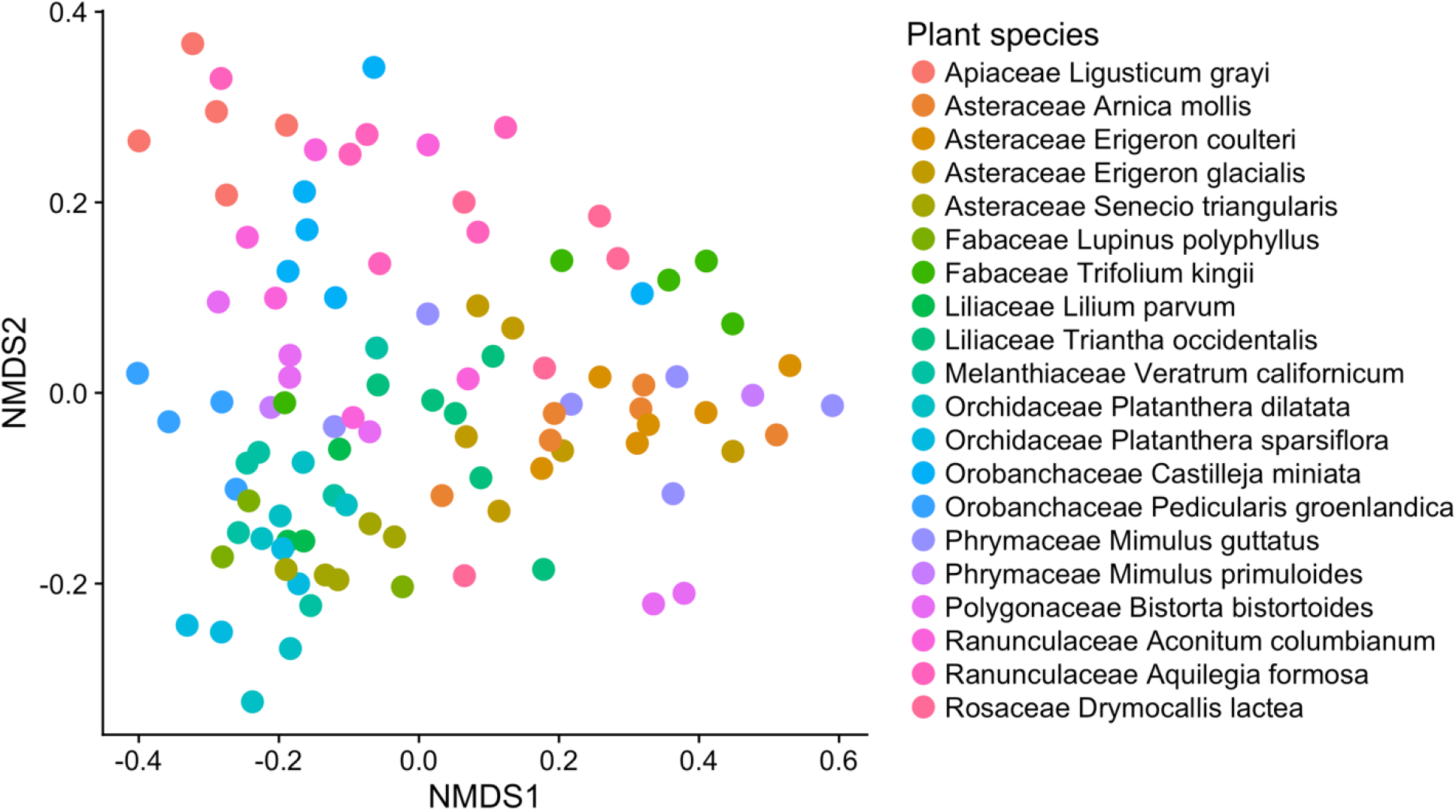
Nonmetric multidimensional scaling plot of floral bacterial communities, with NMDS based on Bray-Curtis dissimilarities, with stress = 0.18. Points are colored by plant species.

At the plant species level, Mantel tests revealed significant correlations between flower visitor dissimilarity and floral bacterial dissimilarity (Fig. 3a; R^2^ = 0.36, p = 0.001). Patterns were consistent and qualitatively similar when either Jaccard or Bray-Curtis dissimilarities were used (Table 1); results based on Jaccard distances are presented below. There was a significant correlation between flower visitor dissimilarity and floral bacterial dissimilarity for each subgroup of flower visitors considered separately, but correlation strength varied by group (Fig. 3b-e). The strongest correlation was between non-bee Hymenoptera and floral bacteria (R^2^ = 0.38, p = 0.001), followed by bees (R^2^ = 0.37, p = 0.001), flies (R^2^ = 0.25, p = 0.002), and beetles (R^2^ = 0.16, p = 0.036).

**Table 1.** Results of Mantel tests for correlations between floral bacterial communities and plant phylogeny and/or flower visitor communities based on dissimilarity matrices using Jaccard or Bray-Curtis distances. Values are R^2^ values and significance levels (* for p ≤ 0.05, ** for p ≤ 0.01, *** for p ≤ 0.005). All correlations were positive.

| **Mantel Test** | **Jaccard** | **Bray-Curtis** |
| --- | --- | --- |
| Plant phylogeny | 0.27 *** | 0.26 *** |
| Whole visitor community | 0.36 *** | 0.38 *** |
| Bee community | 0.37 *** | 0.36 *** |
| Non-bee Hymenoptera community | 0.38 *** | 0.38 *** |
| Fly community | 0.25 ** | 0.27 *** |
| Beetle community | 0.16 * | 0.18 * |
| **Partial Mantel test** | **Jaccard** | **Bray-Curtis** |
| Whole visitor community<br>(controlling for plant phylogeny) | 0.39 *** | 0.41 *** |

**Figure 3.**
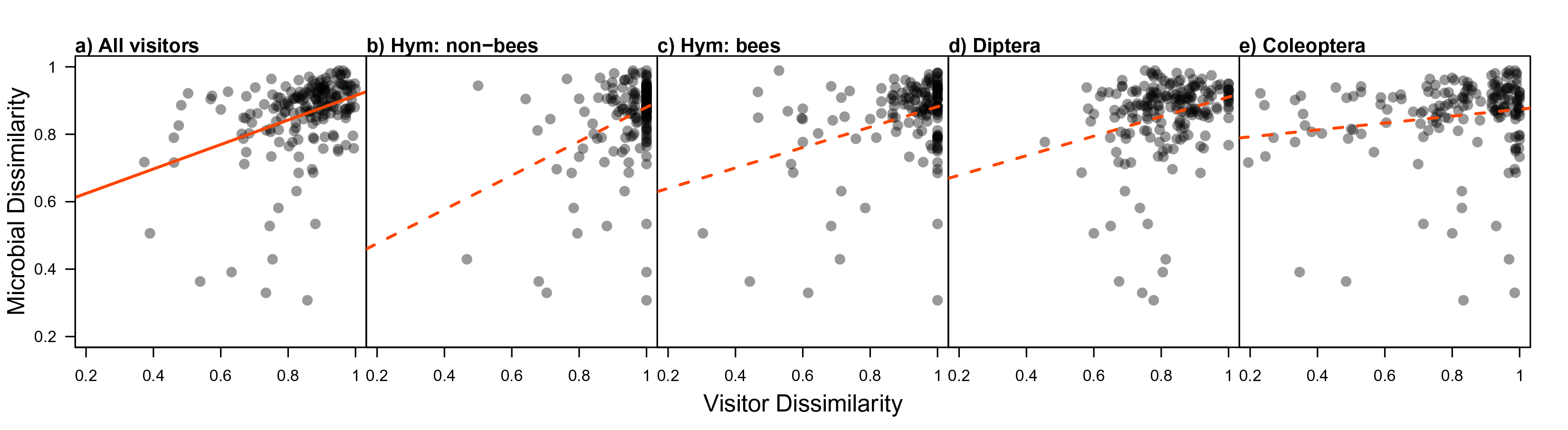
Across plant species, there was a significant correlation between flower visitor and floral bacteria communities for the whole visitor web (a), non-bee hymenopteran visitors (b), bee visitors (c), fly visitors (d) and beetle visitors (e). Points represent dissimilarity values between plant species based on either dissimilarity in microbial community composition and floral visitor community.

Plant phylogenetic distance was also associated with bacterial community dissimilarity (R^2^ = 0.27, p = 0.005); however, even after correcting for plant phylogenetic distance using a partial Mantel test, there remained a strong, significant correlation between flower visitor dissimilarity and floral bacteria dissimilarity (R^2^ = 0.39, p = 0.001; Table 1).

### Flower visitor and floral bacterial network architecture

#### Network macrostructure

The plant – floral bacteria network was significantly more nested than the plant – flower visitor network (Fig. 4a,b). Both networks were composed of only one compartment, and did not have significantly different levels of Modularity Q (Fig. 4c,d). Metrics based on the rarefied plant – floral bacteria matrix did not differ qualitatively from metrics based on the un-rarefied matrix (see caption of Fig. 4).

**Figure 4.**
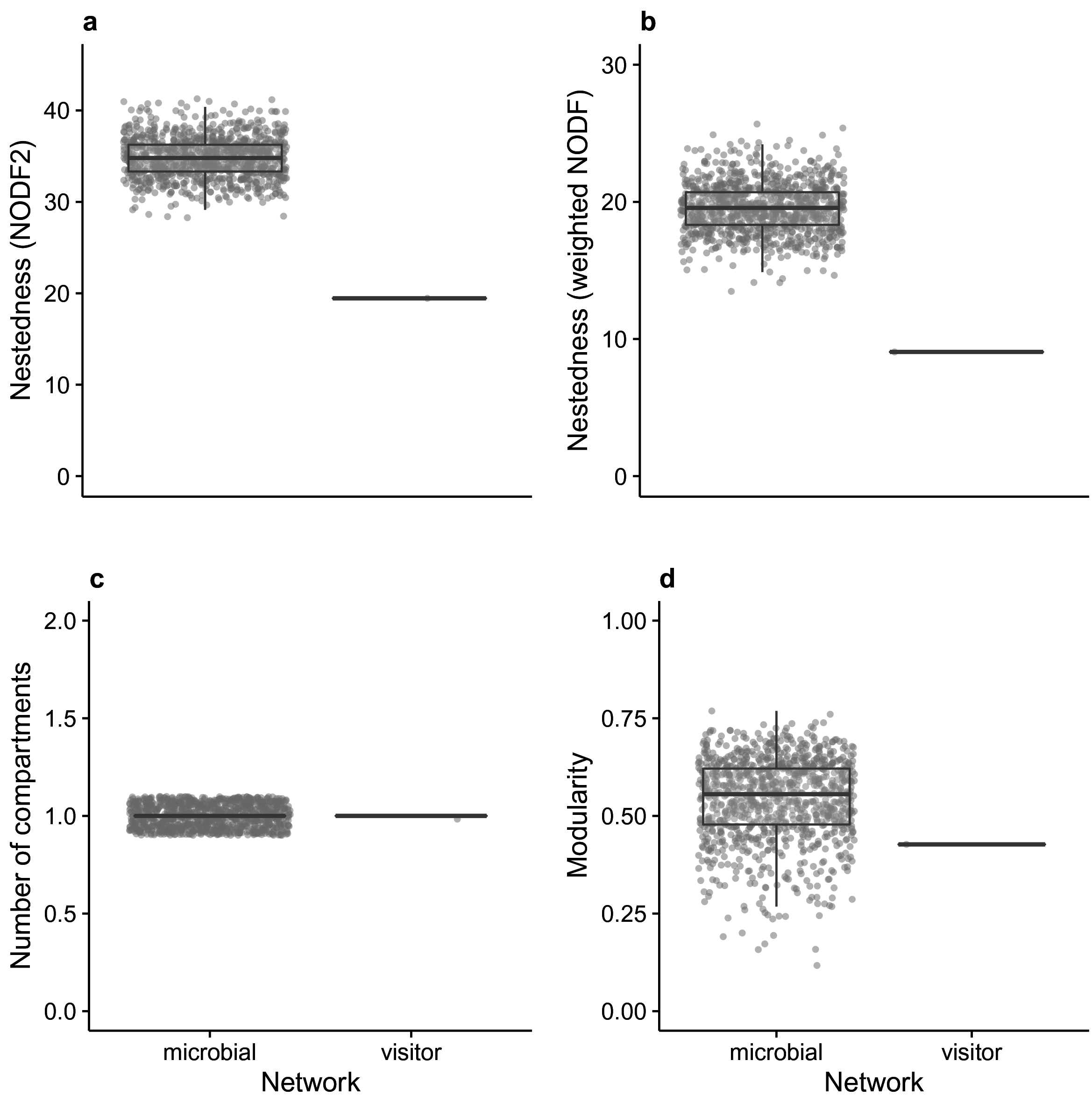
The floral bacteria network was significantly more nested than the flower visitor network considering un-weighted nestedness (a) and weighted nestedness (b). The flower visitor and floral bacteria web had the same number of compartments (c) and average modularity score (d). For the bacterial network, metrics calculated with entire network were similar to the rarefied network: nestedness = 26.63, weighted nestedness = 19.45, number of compartments = 1, and modularity Q = 0.61. Microbial networks were resampled 1000 times and all iterations are shown.

#### Network microstructure

Plant species comprising the nested core of the flower visitor network were not necessarily in the nested core of the floral bacteria network; there was no significant correlation between plant nested rank in both networks (Fig. 5a; R^2^ = 0.04, p = 0.62). Similarly, plants that were hubs of the flower visitor network were not necessarily hubs of the floral bacteria network. There was no significant relationship between plant degree (number of interaction partners) in the visitor and bacterial webs (Fig. 5b; R^2^ = 0.05, p = 0.76). There was no significant relationship between the closeness centrality of plant species in the flower visitor network and floral bacteria network (Fig. 5c; R^2^ = 0.03, p = 0.53).

**Figure 5.**
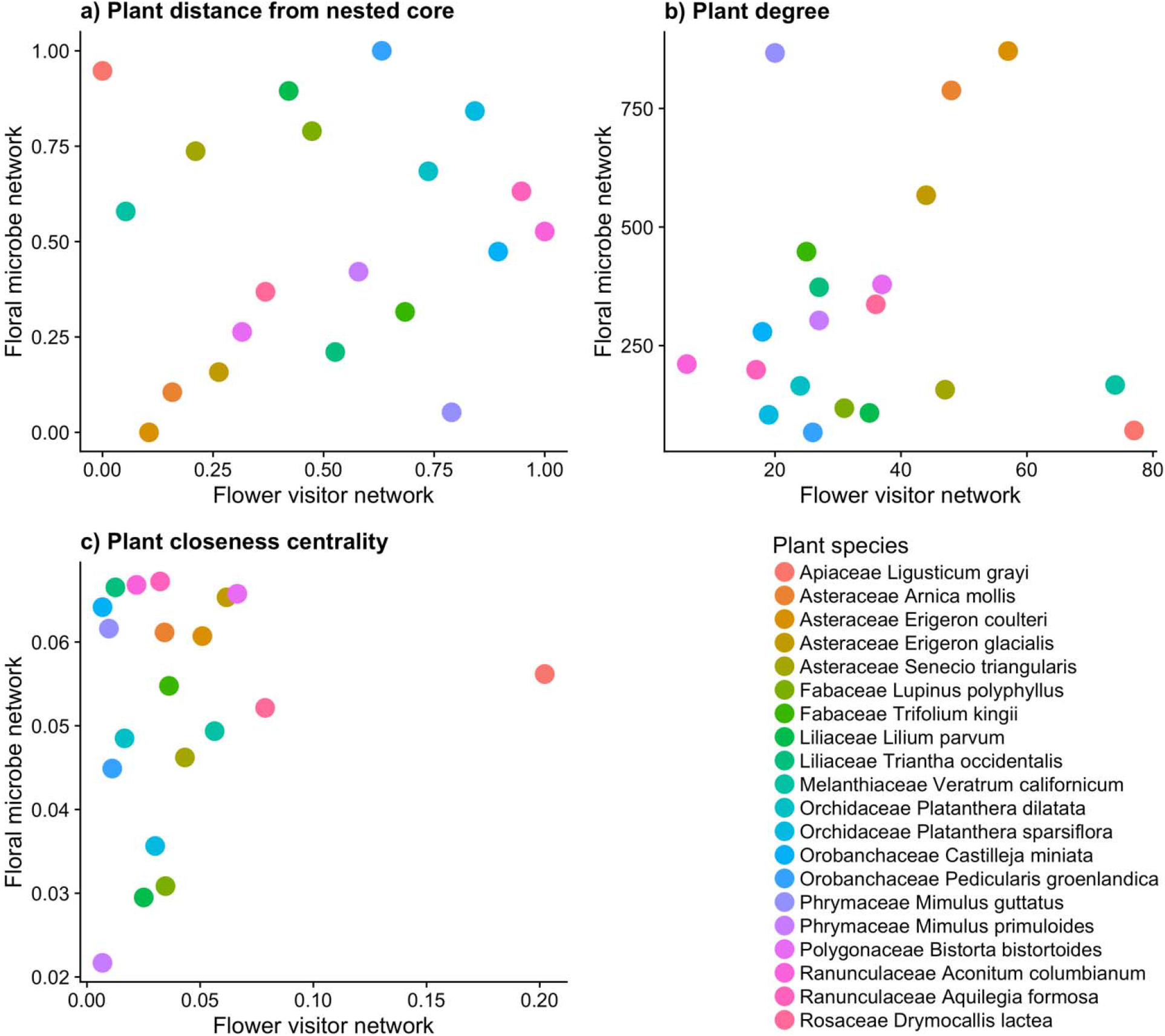
Comparison of plant species network position in flower visitor and floral bacteria networks. (a) Plants in closer proximity to the nested core of the flower visitor network (closer to 0) were not necessarily close to the nested core of the floral bacteria network. (b) Plant species with a high number of flower visitor species (degree) were not necessarily associated with a high number of bacterial OTUs. (c) The closeness centrality of plant species was not related in flower visitor and floral bacteria web. Points represent a single value for each plant species.

## Discussion

The analysis of linked networks presented here reveals that although shared insect visitors of plants likely contribute to microbial community similarity between plant species, additional plant traits influence microbial and insect visitor community networks differently. Our unique network approach revealed distinct network macro and micro-structure features between the linked networks, suggesting that either different plant traits filter floral visitors compared to microbial inhabitants, or that subsequent processes, including environmental filtering, competitive dynamics or plants as bacterial source pools, may influence the similarity of microbial communities among plant species.

Using only analyses of community dissimilarity, a typical approach in microbial community analysis, we would have concluded that dispersal is the major driver of bacterial species composition in floral hosts. However, analyzing these same data using the network approach offered the additional insight that processes other than insect-mediated dispersal are clearly important in structuring floral bacterial communities. For example, *Ligusticum grayi*, an umbel, was the plant species with the greatest degree and closeness centrality in the floral visitor network (Fig. 5), likely due to its accessible morphology. Yet, *L. grayi* was characterized by a low degree and great distance from the nested core in plant-bacterial networks. In contrast, plant species in the Asteraceae (e.g., *Erigeron* spp.) were close to the nested core and showed high closeness centrality in plant-microbe network, but no consistent patterns in the flower visitor network. Taken together, these patterns suggest that despite a likely role for visitor communities in microbial dispersal, distinct processes influence the structure of the floral visitor network compared to bacterial communities on flowers, which may instead be affected by plant volatile composition (Burdon et al. 2018), floral microenvironment, UV exposure, competitive interactions, and historical contingency (Chappell and Fukami 2018). More broadly, we suggest that the linked network approach can offer significant insight into factors that structure microbial communities across hosts that differ both in dispersal frequency and species input, and environmental characteristics compared to the analyses typically applied to microbial community data. Specifically, the use of community similarity or clustering-based algorithms may overlook biologically meaningful features of complex multivariate data, particularly when multilevel network information is available.

Unlike other systems where bacterial dispersal is difficult or near impossible to track, our flower visitor – plant – floral bacteria system allowed us to construct a detailed network of potential microbial vectors to plant hosts. Previous studies have used visitor guilds or observed single plant species to link visitor composition to variation in microbial community structure (Herrera et al. 2009, Vega et al. 2009, Samuni-Blank et al. 2014) but this study offers detailed species-level identification of floral visitors including predators and parasitoids across a relatively diverse coflowering plant community. This rich dataset allowed us to examine the relative contribution of each taxonomic guild of floral visitors. Indeed, a large proportion of the variation in floral bacteria species composition across plant species was explained by the species composition of floral visitors (even when controlling for plant phylogenetic distance; Fig. 3a, Table 1). Further, the correlation strength between visitor and bacterial communities varied when considering different subgroups of the flower visitor community (Fig. 3b-e, Table 1). Interestingly, similar results were found using Bray-Curtis and Jaccard dissimilarity matrices (Table 1), suggesting that both the incidence and abundance of bacteria are influenced by dispersing insects. Past studies of floral microbe communities have tended to focus on only the primary pollinators as microbe vectors (McArt et al. 2014). However, poor vectors of pollen could be just as effective in vectoring microbes as pollinators. Previous work has shown that herbivores (Samuni-Blank et al. 2014) and nectar robbers (Zemenick et al. 2018) have distinct effects on microbial community composition. In the current study, a strong correlation between non-bee Hymenoptera and floral bacterial communities (Fig. 3b,c) suggests that this may hold for floral communities as well. Although the current approach can identify key taxa that effectively link all flower types through microbial dispersal, interactions between visitor and floral morphology that make insect species inconsistent vectors of bacteria (e.g. beetles contact the stigma in some plant species, but not all) may not be detectable. Therefore, future work studying the dispersal or community assembly of floral microbes should not be limited to visitors presumed to be the most efficient pollinators. More generally, this result emphasizes that vector identity is important, but often not considered in metacommunity studies.

The multilevel network framework can be a useful tool to understand the ecology and assembly of linked communities. By building a multilevel plant – flower visitor – floral microbe network as a model system, we show that plant species with similar visitor communities tend to have similar bacterial communities, and that the identity of visitors (disperser) may be more important than dispersal rate in structuring floral bacterial communities. Further, we show that plants occupied divergent positions in the flower visitor and floral bacteria network, suggesting an important role for habitat filtering. Taking these results together, it seems that dispersal is important in determining similarity in microbial communities across plant species, but not as important in determining structural features of plant-floral microbe network. Using a uniquely tractable system to estimate dispersal, we demonstrate that a multilevel network approach is therefore useful in estimating the relative importance of dispersal and other factors of assembly (such as habitat filtering or historical contingency) that cannot be addressed considering networks as separate entities.

## Declarations

The authors thank Philip Campos, Ann Le, Sam Rusoff, Alex Sinclair, and Marshall McMunn for help in the field; Bodil Cass, Adela Contreras, and Griffin Hall provided help in the lab. Help with arthropod identification was graciously provided by: Robbin Thorp, Terry Griswold, Jack Neff, and Jenny VanWyk (bees); Martin Hauser, Chris Borkent, and Stephen Gaimari (Diptera); Brendan Boudinot (Formicidae); Steve Heydon (Parasitic Hymenoptera); Lynn Kimsey (Aculeate Hymenoptera); Moria Robinson (Lepidoptera); Al Wheeler (Hemiptera); and Joel Ledford (Arachnida). Faerthen Felix helped with plant identification. Jeff Brown and Sagehen Creek Field Station were critical to facilitating field work. This work was supported by the National Science Foundation Graduate Research Fellowship DGE-1148897 and Doctoral Dissertation Improvement Grant DEB-1501620 and USDA Hatch Multistate NE1501. Any opinions, findings, and conclusions or recommendations expressed in this material are those of the author(s) and do not necessarily reflect the views of the National Science Foundation.

## References

Aleklett, K., M. Hart, and A. Shade. 2014. The microbial ecology of flowers: an emerging frontier in phyllosphere research. Botany 92:253–266.

Bascompte, J., P. Jordano, C. J. Melián, and J. M. Olesen. 2003. The nested assembly of plant-animal mutualistic networks. Proceedings of the National Academy of Sciences of the United States of America 100:9383–9387.

Belisle, M., K. G. Peay, and T. Fukami. 2012. Flowers as islands: spatial distribution of nectar-inhabiting microfungi among plants of Mimulus aurantiacus, a hummingbird-pollinated shrub. Microbial Ecology 63:711–8.

Block, A. K., E. Yakubova, and J. R. Widhalm. 2019. Specialized naphthoquinones present in Impatiens glandulifera nectaries inhibit the growth of fungal nectar microbes. Plant Direct 3:e00132.

Brysch-Herzberg, M. 2004. Ecology of yeasts in plant-bumblebee mutualism in Central Europe. FEMS microbiology ecology 50:87–100.

Burdon, R. C., R. R. Junker, D. G. Scofield, and A. L. Parachnowitsch. 2018. Bacteria colonising Penstemon digitalis show volatile and tissue-specific responses to a natural concentration range of the floral volatile linalool. Chemoecology 28:11–19.

Burns, A. R., W. Z. Stephens, K. Stagaman, S. Wong, J. F. Rawls, K. Guillemin, and B. J. M. Bohannan. 2016. Contribution of neutral processes to the assembly of gut microbial communities in the zebrafish over host development. ISME Journal 10:655–664.

Carter, C., and R. W. Thornburg. 2004. Is the nectar redox cycle a floral defense against microbial attack? Trends in Plant Science 9:5–9.

Chappell, C. R., and T. Fukami. 2018. Nectar yeasts: a natural microcosm for ecology. Yeast. 35(6):417–423

Deshpande, V., Q. Wang, P. Greenfield, M. Charleston, A. Porras-Alfaro, C. R. Kuske, J. R. Cole, D. J. Midgley, and N. Tran-Dinh. 2016. Fungal identification using a Bayesian classifier and the Warcup training set of internal transcribed spacer sequences. Mycologia 108:1–5.

Dormann, C., and R. Strauss. 2013. Detecting modules in quantitative bipartite networks: the QuaBiMo algorithm. arXiv preprint arXiv 1304.3218.

Fontaine, C., P. R. Guimarães, S. Kéfi, N. Loeuille, J. Memmott, W. H. van der Putten, F. J. F. van Veen, and E. Thébault. 2011. The ecological and evolutionary implications of merging different types of networks. Ecology Letters 14:1170–1181.

Friesen, M. L. 2013. Microbially Mediated Plant Functional Traits. Molecular Microbial Ecology of the Rhizosphere 1:87–102.

Fründ, J., K. S. McCann, and N. M. Williams. 2016. Sampling bias is a challenge for quantifying specialization and network structure: Lessons from a quantitative niche model. Oikos 125:502–513.

Graystock, P., D. Goulson, and W. O. H. Hughes. 2015. Parasites in bloom: flowers aid dispersal and transmission of pollinator parasites within and between bee species. Proceedings of the Royal Society B. 282: 20151371

Herrera, C. M., C. de Vega, A. Canto, and M. I. Pozo. 2009. Yeasts in floral nectar: a quantitative survey. Annals of Botany 103:1415–1423.

Huang, M., A. M. Sanchez-moreiras, C. Abel, R. Sohrabi, S. Lee, J. Gershenzon, and D. Tholl. 2012. The major volatile organic compound emitted from Arabidopsis thaliana flowers, the sesquiterpene (E)-b-caryophyllene, is a defense against a bacterial pathogen. New Phytologist 193:997–1008.

Junker, R. R., T. Romeike, A. Keller, and D. Langen. 2014. Density-dependent negative responses by bumblebees to bacteria isolated from flowers. Apidologie 45:467–477.

Liaw, A. and M. Wiener (2002). Classification and Regression by randomForest. R News 2(3), 18–22.

McArt, S. H., H. Koch, R. E. Irwin, and L. S. Adler. 2014. Arranging the bouquet of disease: Floral traits and the transmission of plant and animal pathogens. Ecology Letters 17:624–636.

McFrederick, Q. S., J. M. Thomas, J. L. Neff, H. Q. Vuong, K. A. Russell, A. R. Hale, and U. G. Mueller. 2017. Flowers and Wild Megachilid Bees Share Microbes. Microbial Ecology 73:188–200.

McMurdie, P. J., and S. Holmes. 2013. phyloseq: An R Package for Reproducible Interactive Analysis and Graphics of Microbiome Census Data. PLoS ONE 8(4):e61217

Ngugi, H. K., and H. Scherm. 2006. Biology of flower-infecting fungi. Annual review of Phytopathology 44:261–82.

Nielsen, A., and J. Bascompte. 2007. Ecological networks, nestedness and sampling effort. Journal of Ecology 95:1134–1141.

Oksanen, J., F.G. Blanchet, M. Friendly, R. Kindt, P. Legendre, D. McGlinn, P.R. Minchin, R. B. O’Hara, G.L. Simpson, P. Solymos, M.H.H. Stevens, E. Szoecs and H. Wagner. 2017. vegan: Community Ecology Package. R package version 2.4-3. https://CRAN.Rproject.org/package=vegan

Peay, K. G., M. G. Schubert, N. H. Nguyen, and T. D. Bruns. 2012. Measuring ectomycorrhizal fungal dispersal: macroecological patterns driven by microscopic propagules. Molecular Ecology 21:4122–4136.

Pilosof, S., M. A. Porter, M. Pascual, and S. Kéfi. 2017. The multilayer nature of ecological networks. Nature Ecology & Evolution 1:1–9.

R Core Team (2017). R: A language and environment for statistical computing. R Foundation for StatisticalComputing, Vienna, Austria. URL https://www.R-project.org/.

Rering, C. C., J. J. Beck, G. W. Hall, M. M. McCartney, and R. L. Vannette. 2017. Nectar-inhabiting microorganisms influence nectar volatile composition and attractiveness to a generalist pollinator. New Phytologist: 220(3):750–759

Samuni-Blank, M., I. Izhaki, S. Laviad, A. Bar-Massada, Y. Gerchman, and M. Halpern. 2014. The role of abiotic environmental conditions and herbivory in shaping bacterial community composition in floral nectar. PloS one 9:e99107.

Shade, A., P. S. McManus, and J. Handelsman. 2013. Unexpected diversity during community succession in the apple flower microbiome. mBio 4:1–12.

Silk, M. J., D. P. Croft, R. J. Delahay, D. J. Hodgson, M. Boots, N. Weber, and R. A. McDonald. 2017. Using Social Network Measures in Wildlife Disease Ecology, Epidemiology, and Management. BioScience 67:245–257.

Stein, R. A. 2011. Super-spreaders in infectious diseases. International Journal of Infectious Diseases 15:e510–e513.

Sugio, A., G. Dubreuil, D. Giron, and J. C. Simon. 2015. Plant-insect interactions under bacterial influence: Ecological implications and underlying mechanisms. Journal of Experimental Botany 66:467–478.

Tylianakis, J. M., E. Laliberté, A. Nielsen, and J. Bascompte. 2010. Conservation of species interaction networks. Biological Conservation 143:2270–2279.

Ushio, M., E. Yamasaki, H. Takasu, A. J. Nagano, S. Fujinaga, M. N. Honjo, M. Ikemoto, S. Sakai, and H. Kudoh. 2015. Microbial communities on flower surfaces act as signatures of pollinator. Scientific Reports 5:1–7.

Vannette, R. L., M.-P. L. Gauthier, and T. Fukami. 2013. Nectar bacteria, but not yeast, weaken a plant-pollinator mutualism. Proceedings of the Royal Society B: Biological Sciences 280:1–7.

de Vega, C., C. M. Herrera, and S. D. Johnson. 2009. Yeasts in floral nectar of some South African plants: Quantification and associations with pollinator type and sugar concentration. South African Journal of Botany 75:798–806.

Venkataraman, A., C. M. Bassis, J. M. Beck, V. B. Young, J. L. Curtis, G. B. Huffnagle, and T. M. Schmidt. 2015. Application of a neutral community model to assess structuring of the human lung microbiome. mBio 6. 1) e02284–14

Wang, Q., G. M. Garrity, J. M. Tiedje, and J. R. Cole. 2007. Naïve Bayesian classifier for rapid assignment of rRNA sequences into the new bacterial taxonomy. Applied and Environmental Microbiology 73:5261–5267.

Zemenick, A. T., J. A. Rosenheim, and R. L. Vannette. 2018. Legitimate visitors and nectar robbers of Aquilegia formosa have different effects on nectar bacterial communities. Ecosphere 9:e02459.

