## Supplementary Information for "Linked networks reveal dual roles of insect dispersal and species sorting for bacterial communities in flowers"

**Figure S1**

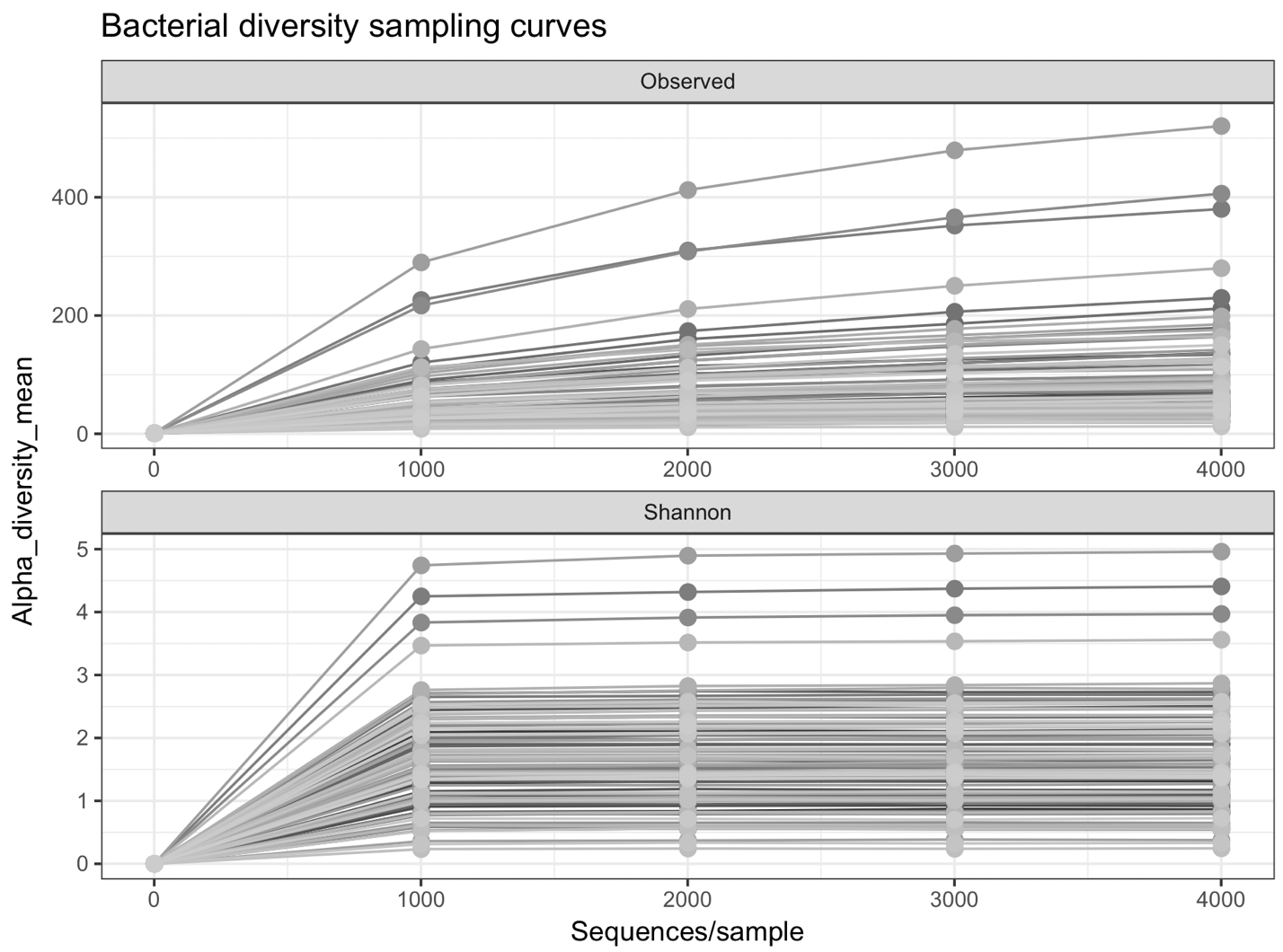

Floral bacteria alpha diversity sampling curves. Rarefying to an even depth of 4,206 reads sufficiently captures bacterial diversity in terms of richness (top panel) and Shannon diversity (bottom panel) within samples since the majority of samples reach an asymptote by 4000 reads for both measures.

**Figure S2**

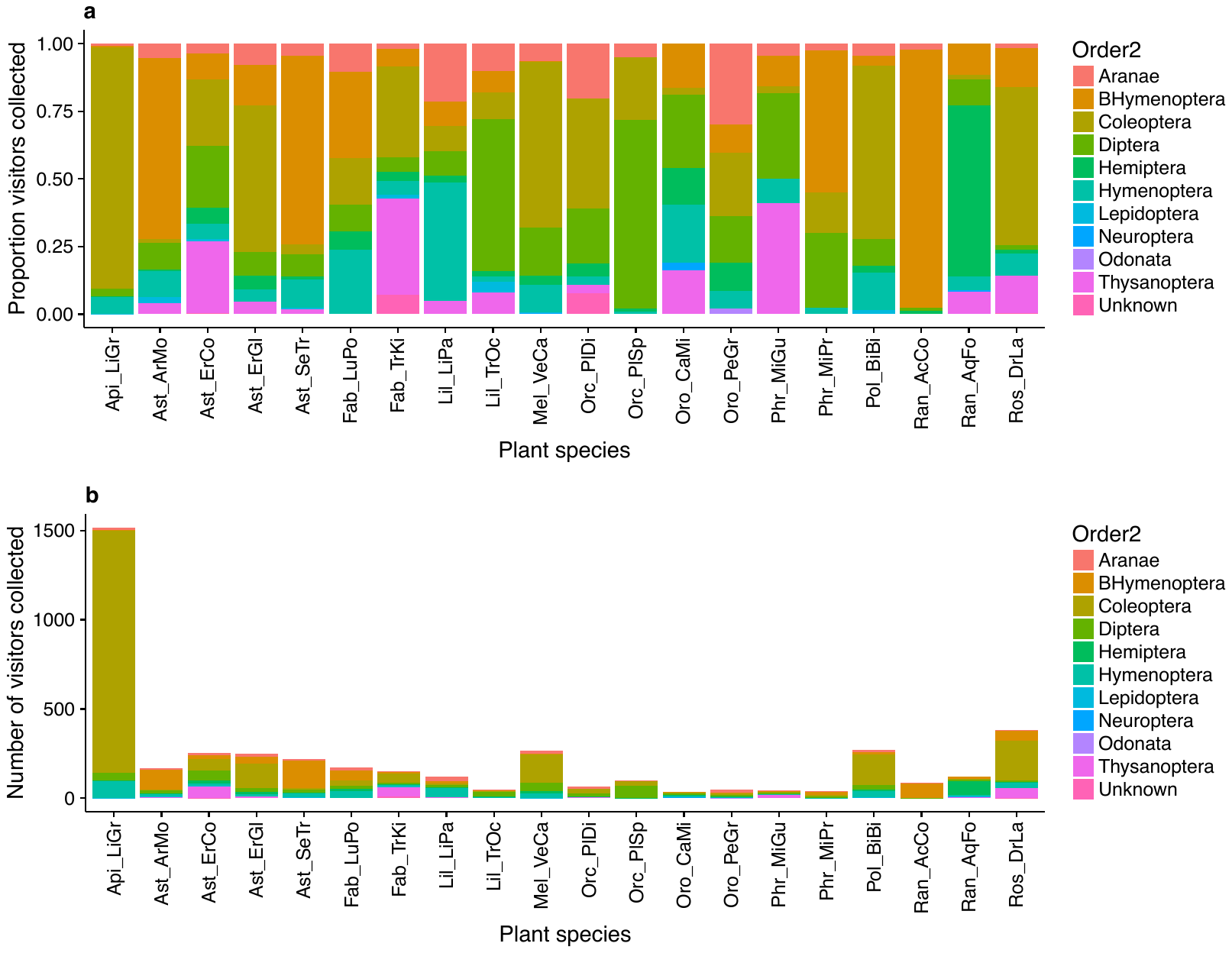

Visitor composition across plant species in terms of (a) the relative abundance and (b) the abundance of arthropod visitors by order. The relative and total abundances are based from the total number of specimens collected from each plant species during 7 hours of observation per species. Note that the Order Hymenoptera was split to show the difference between bees (“BHymenoptera”) in orange and non-bee Hymenoptera in teal.

**Figure S3**

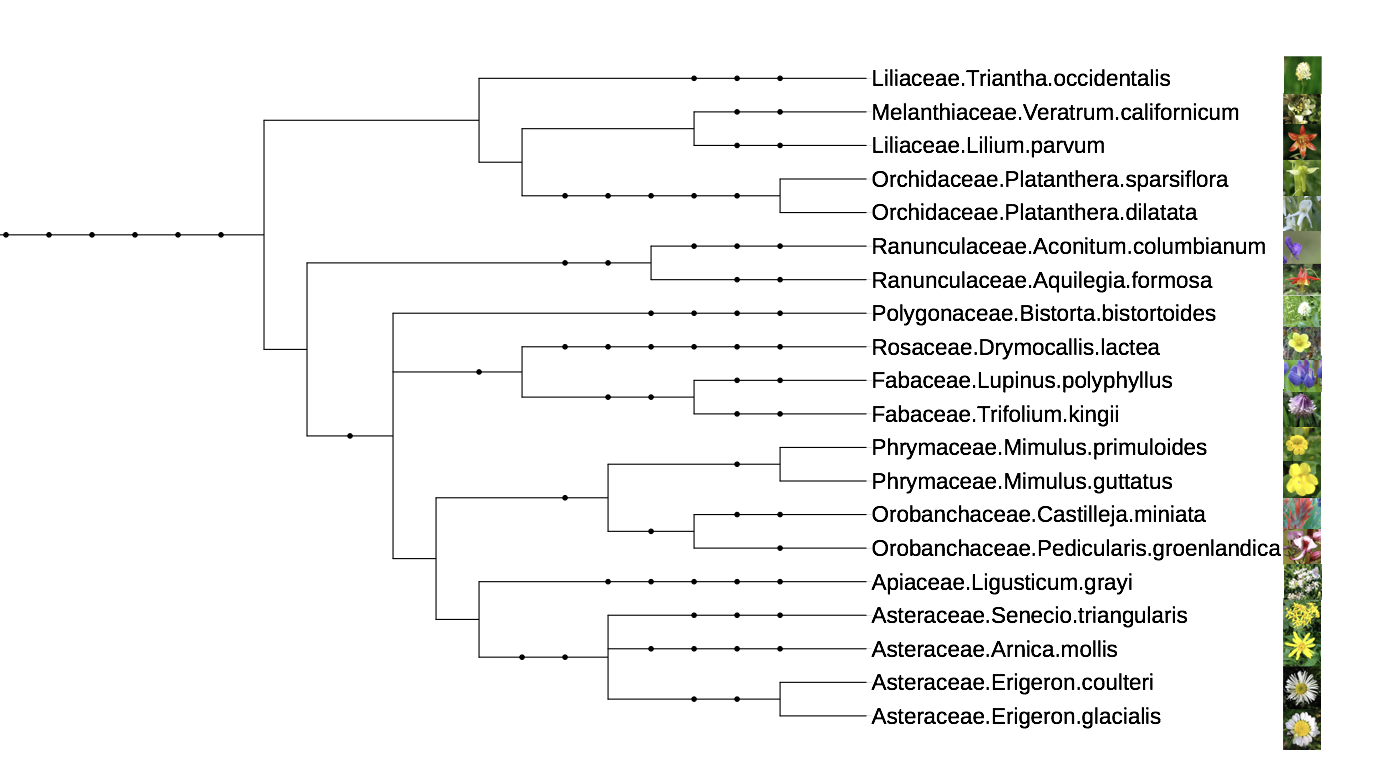

Phylogeny of focal plant species made with phyloT: A phylogenetic tree generator, based on NCBI taxonomy (phylot.biobyte.de/).

**Figure S4**

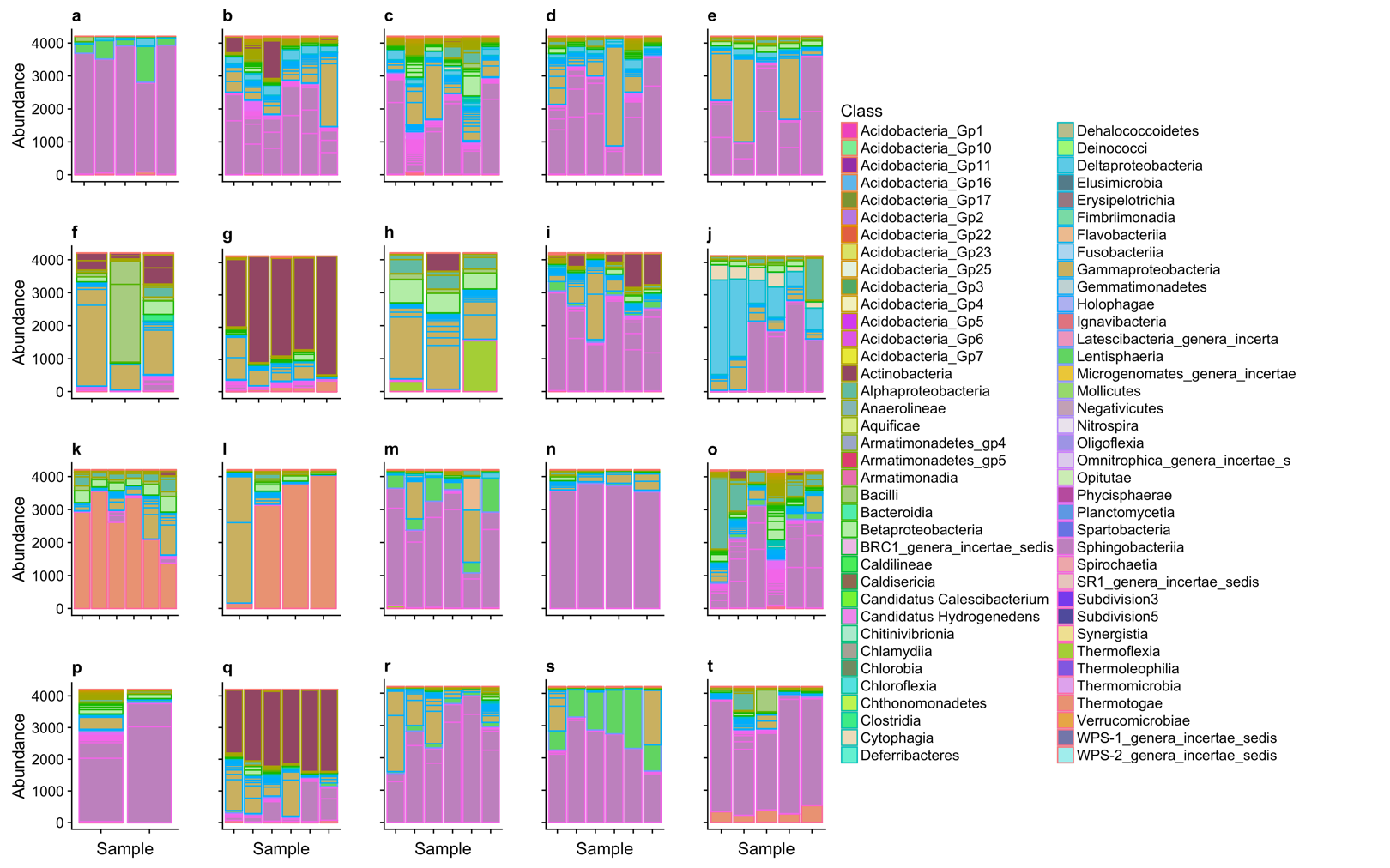

Relative abundance (number of reads after rarefaction) of bacterial classes isolated from flower species in the family Apiaceae (a), Asteraceae (b-e), Fabaceae (f-g), Liliaceae (h-i), Melanthiaceae (j), Orchidaceae (k-l), Orobanchaceae (m-n), Phrymaceae (o-p), Polygonaceae (q), Ranunculaceae (r-s), and Roseaceae (t). The most common being Actinobacteria, Alphaproteobacteria, Bacili, Betaproteobacteria, Cytophaga, Deltaproteobacteria, Flavobacteria, Gammaproteobacteria, Lentisphaeria, Sphingobacteria, and Thermotogae. Individual bars represent samples processed each with five flowers. Panels correspond to the 20 focal plant species: a) *Lingusticum grayi*, b) *Arnica mollis*, c) *Erigeron coulteri,* d) *E. glacialis,* e) *Senecio triangularis*, f) *Lupinus polyphyllus,* g) *Trifolium kingii,* h) *Lilium parvum,* i) *Triantha occidentalis*, j) *Veratrum californicum*, k) *Platanthera dilatata,* l) *P. sparsiflora*, m) *Castilleja miniata,* n) *Pedicularis groenlandica,* o) *Mimulus guttatus,* p) *M. primuloides,* q) *Bistorta bistortoides,* r) *Aconitum columbianum,* s) *Aquilegia* *formosa* and t) *Drymocallis lactea.*

**Figure S5**

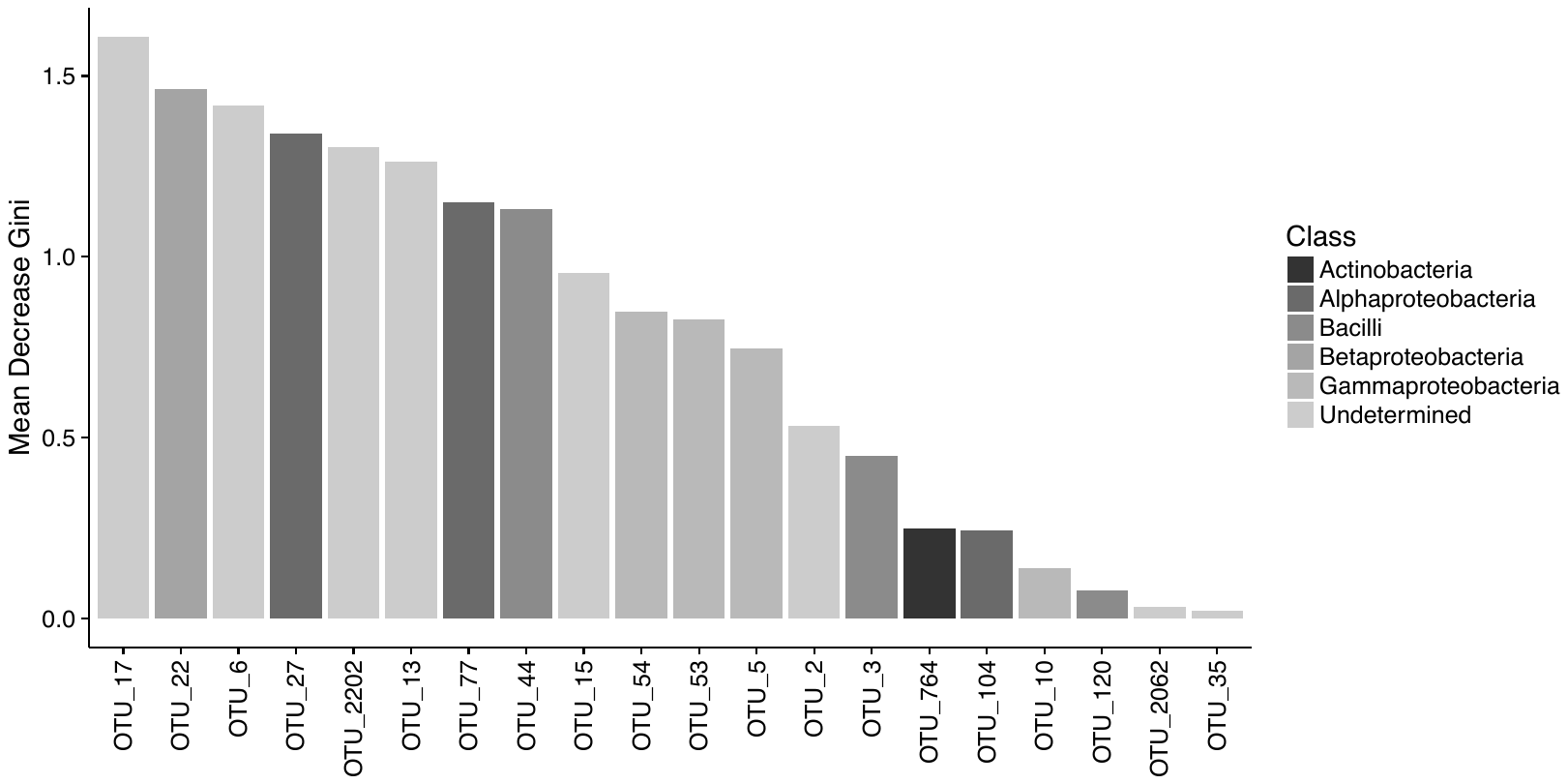

The 20 most influential bacterial OTUs in splitting floral bacterial OTUs across plant species as determined by random forest. One OTU was assigned to the Class Actinobacteria (OTU 764), three were assigned to Alphaproteobacteria (OTUs 27, 77, 104), three to Bacilli (OTUs 3, 44, 120), one to Betaproteobacteria (OTU 22), four to Gammaproteobacteria (OTUs 5, 10, 53, 54), and eight were not assigned to Class with high confidence (OTUs 2, 6, 13, 15, 17, 35, 2062, and 2202). See Table S2 for taxonomic assignments of OTUs.

**Table S1**

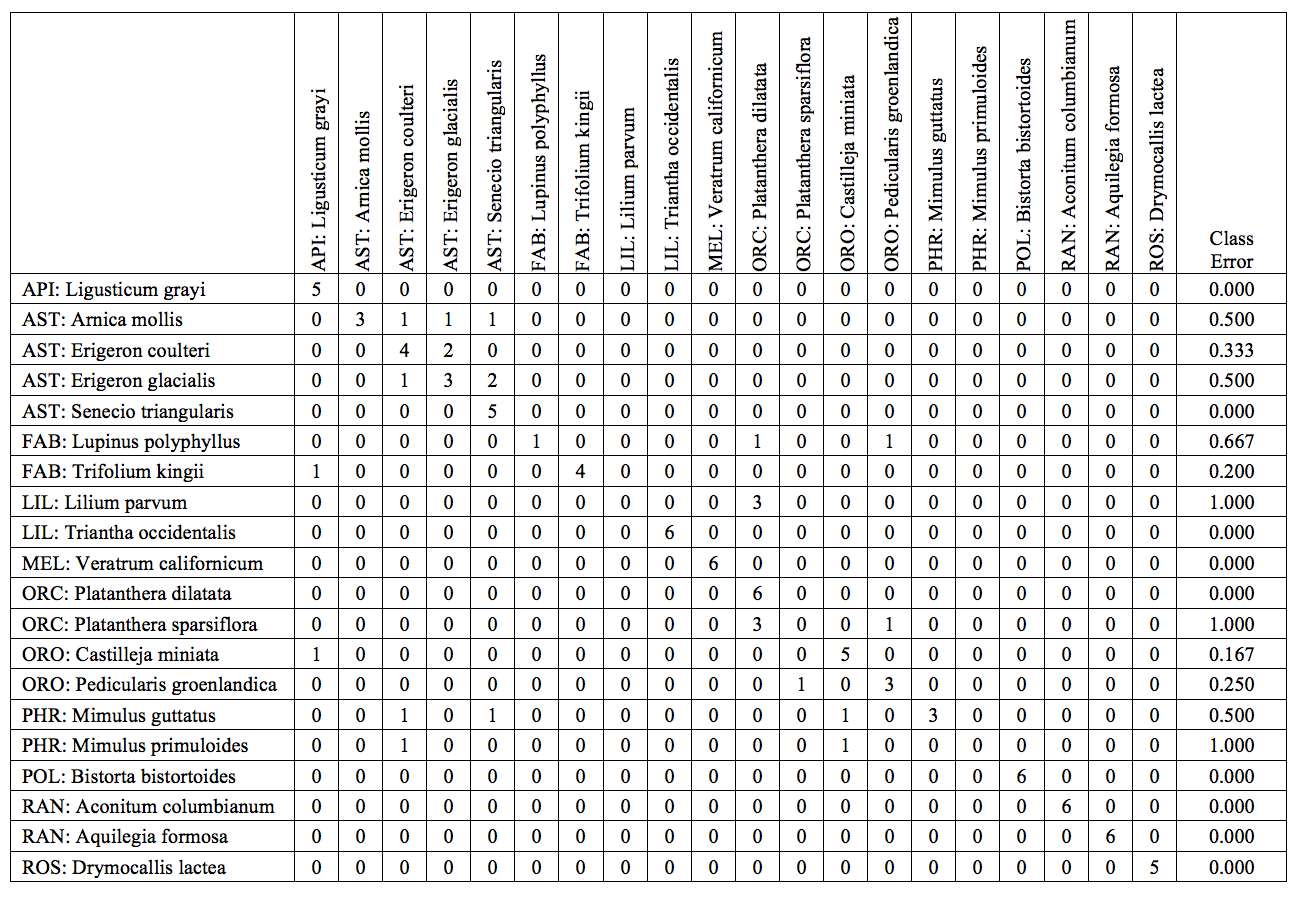

Confusion matrix from random forest identifying OTUs important in splitting microbial communities across plant species.

**Table S2**

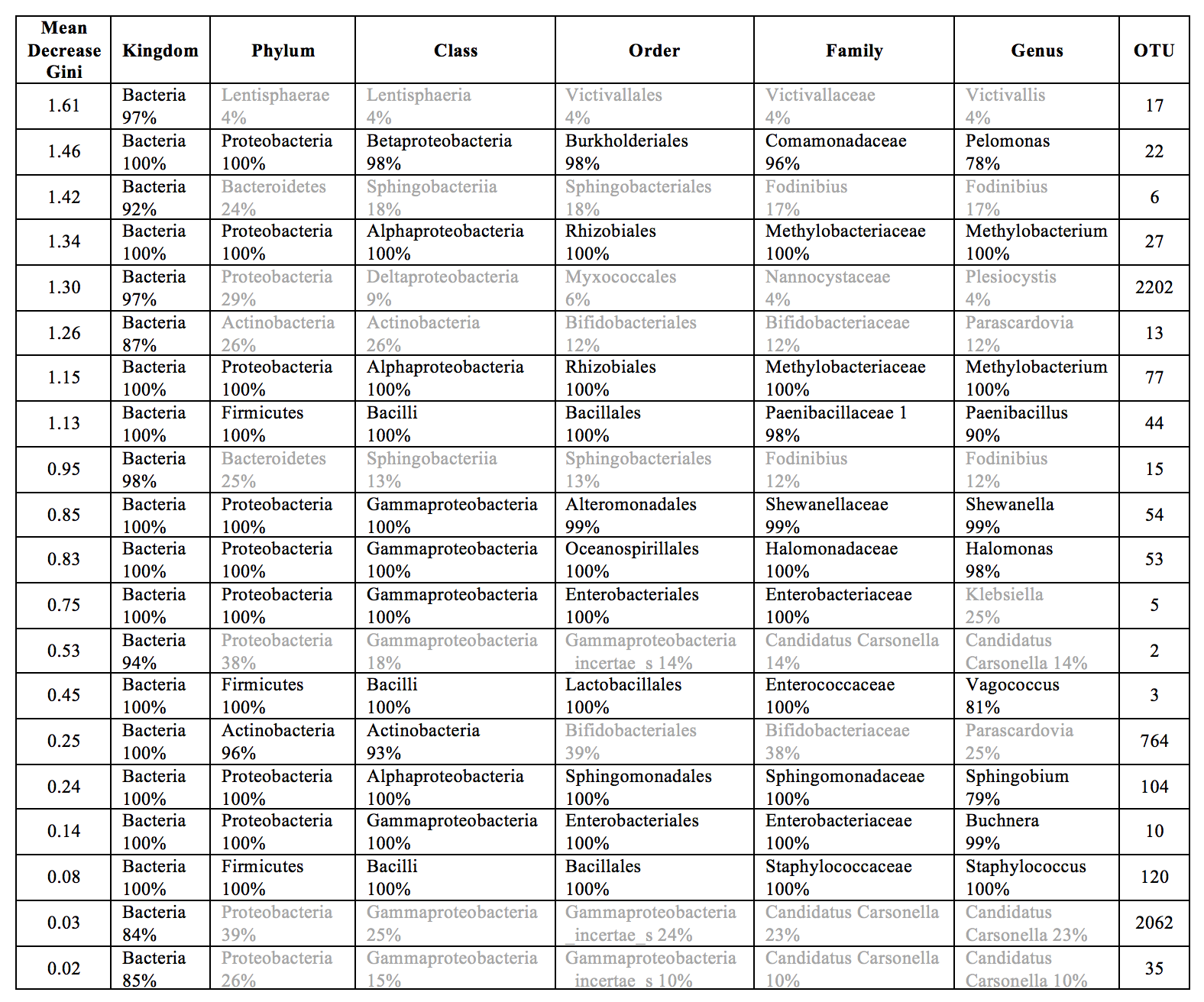

Taxonomic classification of the 20 most influential OTUs in driving differences across plant species. Percentages refer to confidence in taxonomic assignments, and text in grey indicates taxonomic assignment ≤ 50%.
